# Neural Oscillatory Signatures of Predictive Processing in Visual Statistical Learning

**DOI:** 10.64898/2026.02.25.707965

**Authors:** Martina Pasqualetti, Jakob C. B. Schwenk, Andrea Alamia

**Author notes:** Corresponding author: Martina Pasqualetti.

## Abstract

Several studies suggest that neural oscillations play a role in cognition, including predictive processing. Recent models propose that alpha-band traveling waves reflect prediction (top-down, frontal to occipital), while bottom-up waves reflect prediction errors. We tested this hypothesis using a visual statistical learning task in which participants (N=31) detected target shapes presented with varying levels of predictability. EEG and pupillometry were recorded throughout the task. Behavioral results showed faster responses for predictable targets, and pupil dilation increased for unexpected shapes. ERP analyses revealed predictability effects on the P300 component across occipital, central, and frontal electrodes, along with modulation of alpha oscillations. However, traveling-wave analyses did not show a clear effect of predictability between conditions. Instead, wave patterns varied with participants’ cognitive strategies: individuals who relied more on statistical regularities showed stronger top-down alpha-band waves. These findings suggest that alpha traveling waves may reflect general cognitive strategies rather than specific predictive processes.

## Introduction

The brain can be interpreted as a predictive machine, capable of identifying statistical regularities to optimize interactions with the environment (Schapiro et al. 2013; Batterink, Paller, and Reber 2019; Conway 2020). According to the Bayesian brain theory, our brain constantly attempts to infer the causes of the surroundings by continuously updating an internal model of the world, thereby generating probabilistic beliefs about the future (Clark 2013; Colombo and Seriès 2012; Doya et al. 2007; Friston 2012; Knill and Pouget 2004; Little and Sommer 2013). Specifically, hierarchically higher brain areas generate predictions about the causes of sensory input. At the same time, lower regions inform the update of such predictions via Prediction Errors (PE) (i.e., the difference between top-down predictions and the actual activity) (Rao and Ballard 1999).

Statistical Learning (SL) is a widely used paradigm for exploring these mechanisms, and it involves the learning of statistical regularities, such as within a sequence of items (Saffran, Aslin, and Newport 1996; Fiser and Aslin 2001; Turk-Browne, Jungé, and Scholl 2005; Schapiro and Turk-Browne 2015; Fiser and Lengyel 2022). Several neurophysiological correlates have been proposed as possible markers of predictive processes, including evoked response potentials (ERPs) (Abla and Okanoya 2009; Fiser and Aslin 2001; Jost et al. 2011), hemodynamic responses (Turk-Browne et al. 2010), functional connectivity (FC) (Rowchan et al. 2025; Tóth et al. 2017), and neural oscillations (Arnal and Giraud 2012; Engel, Fries, and Singer 2001; Roehe et al. 2021; Sáringer et al. 2023; Zhou et al. 2020).

Regarding neural oscillations, an increasing number of studies are also considering their spatial component, specifically how they propagate across brain regions as traveling waves (TW) (Muller et al. 2018a). In scalp electrophysiological (EEG) recordings, recent studies have associated TW propagation with distinct cognitive phenomena, such as visual perception (Lozano-Soldevilla and VanRullen 2019; Pang (庞兆阳), Alamia, and VanRullen 2020), working memory (Zeng, Sauseng, and Alamia 2024), as well as motor excitability (Haigh et al. 2025).

Interestingly, recent work related alpha-band (8-12 Hz) oscillations and TWs to predictive processes (Alamia and VanRullen 2019; Friston 2019). For example, previous experimental work has shown that pre-stimulus alpha oscillations are enhanced for predicted visual stimuli compared to unpredicted ones, suggesting that alpha oscillations may carry top-down sensory predictions (Mayer et al. 2016; Roehe et al. 2021). Other lines of work have also related alpha oscillations to temporal predictability (Arnal and Giraud 2012). Further computational simulations suggested that oscillatory alpha-band TW may increase due to cortico-cortical interactions driven by predictive coding mechanisms (Alamia and VanRullen 2019; Schwenk and Alamia 2024). These models suggest that forward (FW) waves propagating from occipital to frontal areas encode sensory information and PEs. In contrast, backward (BW) waves, spreading from frontal to occipital regions, are believed to carry top-down predictions. Despite recent work addressing the relationship between alpha-band oscillations and predictive coding (Alamia et al. 2020, 2025; Arnal and Giraud 2012; Friston 2019), clear experimental evidence for this link remains lacking.

Here, we assess this compelling hypothesis using an SL paradigm while measuring EEG and pupil responses. Participants performed a visual detection task, in which they identified a target shape embedded in a sequence of geometric stimuli. Conditional probabilities were manipulated to systematically vary sequence predictability across multiple levels, spanning from highly predictable to entirely unpredictable. Based on the literature, we hypothesized that BW waves, reflecting top-down predictions, would increase with more predictable sequences. In addition, our method of controlling sequence predictability enabled us to define unexpected events, such that violations became increasingly surprising as predictability increased. This approach allowed us to examine whether FW waves reflect PEs and, specifically whether these responses are modulated by the degree of unpredictability associated with the transition. Statistical learning was assessed behaviorally using reaction times as a function of target transition predictability. In addition, ERPs and pupillary responses time-locked to shape onset were quantified for each condition. Cortical TWs were also quantified using two distinct and complementary analytical approaches. Overall, we present a comprehensive description of behavioral and electrophysiological measures to characterize the correlates of visual statistical learning.

## Materials and methods

### Participants

Thirty-two healthy young adults participated in this study; all of them had normal or corrected-to-normal vision. One of them had to be removed from the analysis due to poor behavioral performance (more than 30% of missed targets, compared to the group mean of 2.95%). The following analyses are based on 31 participants (23 males, mean age=26.2, SD=2.90, range=22-33). All participants gave informed consent and were reimbursed for their time. All procedures were approved by the local ethics committee ‘Comité de protection des Personnes Sud Méditerranée 1’ (ethics approval number 2022-A02453-40), adhered to the guidelines for research at the ‘Centre de Recherche Cerveau et Cognition’, and were conducted according to ethical standards of the 1964 Helsinki Declaration.

### Apparatus

#### The experiment took place in a dark and quiet room

All tests were conducted on a PC (Precision 5820 Tower, Dell Inc., Round Rock, Texas, USA) with an NVIDIA graphics card (T1000 8GB, Nvidia Corporation, Santa Clara, California, USA). Visual stimuli were generated and controlled using MATLAB (R2019b, The MathWorks Inc., Natick, MA) and PsychToolbox (Brainard 1997; Kleiner et al. 2007; Pelli 1997).

All the stimuli were displayed on a monitor (Display++ LCD Monitor, 1440x1080 pixels, 100Hz refresh rate, Cambridge Research Systems, United Kingdom). For the entire duration of the experiment, participants had their heads stabilized by a chin rest positioned 57cm from the screen. Before each session, participants could adjust the chin rest to improve their posture. Continuous brain activity was recorded from the subjects using a 64-channel active BioSemi EEG system, at a sampling rate of 2048Hz. The pupil diameter was measured using an EyeLink 1000 Plus video-based eye tracker (SR Research), which recorded monocular pupil size (in arbitrary units) and eye movements at 1000Hz. Calibration of the eye tracker was performed once at the beginning of the experiment or every time the participant adjusted the chin rest.

### Task and Stimuli

Participants performed a visual detection task in which four white geometrical shapes (square, circle, triangle, and diamond) were presented on a black screen, one at a time, in a continuous stream at a 1Hz temporal frequency (Figure 1A). The four shapes had a radius of 3 degrees and a thickness of 0.5 degrees of visual angle. Shapes appeared centrally, slightly above the fixation cross (0.66 degrees of visual angles from the center of the shape), which was white and centered, with a radius of 0.33 degrees of visual angle.

**Figure 1.**
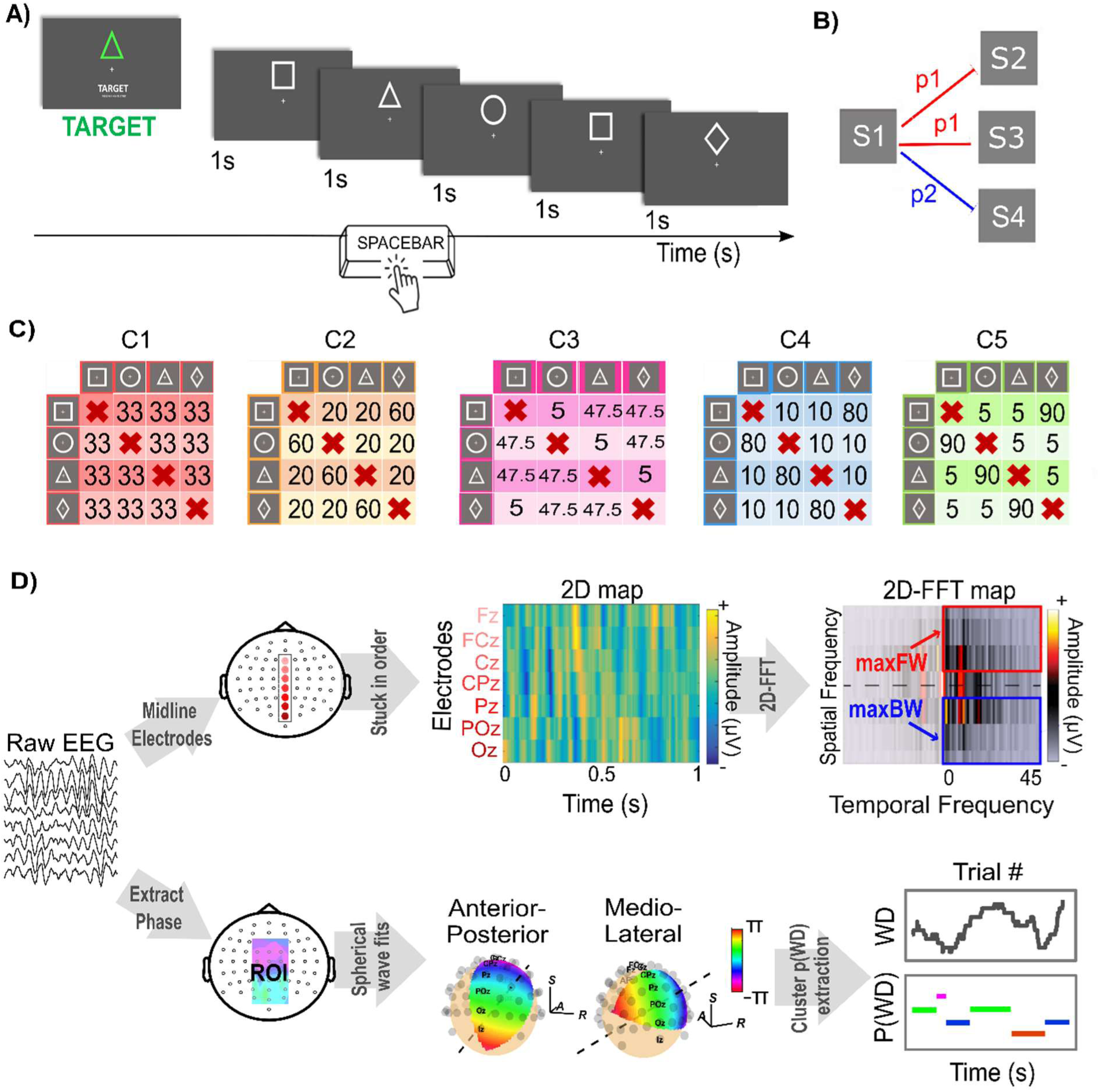
Experimental design and wave quantification methods. A) Trial design. Each trial begins with the presentation of the Target, which is shown here as a green triangle. Once the sequence of shapes begins, each shape lasts 1 second. Subjects were instructed to respond as fast as possible when detecting the target. B) The sequence followed a probabilistic Markovian process, where each shape can transition to another shape with a certain probability defined by the experimental condition. C) Probability tables for all five conditions arranged in order of increasing predictability. Each table shows the transition probability (in percent) between any two shapes. For example, for the rightmost one, after the square, there is a 5% probability of a circle or triangle, and a 90% probability of a diamond. The circle and triangle shapes are considered “rar” transitions, while the diamond is the “expected” one. D) TW quantification methods. Upper panels: 2D-FFT. The raw EEG signals are extracted from the seven midline electrodes (Fz, FCz, Cz, CPz, Pz, POz, Oz) and stacked to obtain 2D maps. After applying a 2D-FFT, we obtain a spectrum with temporal and spatial frequencies on the x- and y-axes respectively. The maximum values in the upper and lower quadrants represent the strength of forward and backward waves, respectively. Lower panel: A spherical wave fit is applied to the alpha-band phase map within a fixed ROI (after applying a power-threshold). For each time-point, the best fit is selected to quantify wave direction and spatial frequency.

The experiment consisted of 15 blocks (∼4min each), each comprising 5 trials. Each trial consisted of a 36-shapes sequence, which the participants initiated by pressing a button. Before each trial, one of the four shapes was designated as the target for that specific trial. Participants were instructed to press the spacebar as fast as possible whenever the target shape appeared. Short breaks were allowed between blocks, and the full experiment lasted about 90min.

At the end of each block, participants received point-based feedback on their RT: they earned or lost 1 point depending on whether their RT was faster/slower than in the previous block. This feedback was added to further incentivize fast responses that rely more strongly on the learned regularity. Lastly, at the end of each block, participants were given a questionnaire in which they were asked to rate, on a scale of 1 to 10, the overall perceived predictability of the stream of shapes in that block.

### Experimental design and procedure

The sequence of shapes within each trial followed a probabilistic Markovian process (Figure 1B), where each shape transitions to one of the three other shapes with different probabilities. We manipulated the predictability of the sequence across five conditions (Figure 1C), defining rare and expected transitions (Table 1). In the four conditions with non-random transitions (C2 to C5), expected transitions have probabilities of 0.9, 0.8, 0.6, and 0.475, while rare transitions have probabilities of 0.05, 0.1, and 0.2, respectively. Note that condition C3 in the table had a slightly different structure in which two expected transitions had a probability of 0.475, and one rare transition had a probability of 0.05. Lastly, one condition (i.e., C1 in the table) was completely unpredictable, with all the transitions having the same probability.

**Table 1.** Probabilities of the shape transitions and entropy for each experimental condition. From condition 1 to condition 5, the reliability of the sequence increases, as indicated by the decrease in entropy, which is measured as − ∑^N^_i_ *p*(*x*_i_) ∗ *log*_2_(*p*(*x*_i_)), where *p*(*x*_i_) indicates the transition probabilities.

|  | **C1** | **C2** | **C3** | **C4** | **C5** |
| --- | --- | --- | --- | --- | --- |
| *Rare event probability* | 33% | 20% | 5% | 10% | 5% |
| *Expected event probability* |  | 60% | 47.5% | 80% | 90% |
| *Entropy* | 1.097 | 0.950 | 0.859 | 0.639 | 0.394 |

The transition matrix remained constant across the 5 trials within each block. Throughout the entire experiment, three blocks per condition were presented in a different randomized order to each subject, with 180 shapes per block (540 for each condition). Target shape varied pseudo-randomly across trials in order to maintain attention. The subdivision of trials was intended solely to allow participants to take short breaks and to change the target.

Before the experiment began, participants were informed that difficulty would vary between blocks, without however being given specific instructions about the specifics of the probabilistic rules, and instructed to respond to the target as quickly as possible.

### Data Analysis

#### Behavioral analysis

During the experiment, we recorded participants’ RTs, their performances (hit rate, miss rate, and false alarm rate), and the score regarding the perceived predictability of the block sequence. Importantly, we also considered answers provided up to 200ms before shape onset as hits or false alarms (on average, across all 31 participants, 2.7% of the targets were anticipated). For these target detections, we hence considered negative RTs.

For all measures except RT, we performed repeated-measures ANOVAs, considering CONDITION as a fixed factor. For the RT, we performed repeated-measures ANOVAs considering CONDITION and SHAPE PROBABILITY (i.e., rare versus expected shape) as fixed factors.

In addition, we defined a Learning Index (LI) to assess the extent to which participants used or did not use the statistical regularities. To do so, we calculated, for each condition separately (except C1, in which there was no regularity, hence no rare or expected events), the *LI*_cond_ as follows:

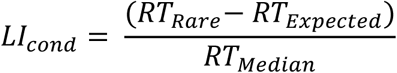

Where *RT*_Rare_and *RT*_Expected_ are the median RT for the rare and expected targets, and *RT*_Median_ is the median RT over all target shapes. To assess the effect of condition on this index, we performed repeated-measures ANOVAs, with CONDITION as a fixed factor.

For each participant, we also calculated the LI across conditions as follows:

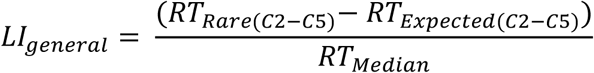

Where *RT*_Rare(C2-C5)_and *RT*_Expected(C2-C5)_are the median RT across conditions C2-C5 for the rare and expected targets, and *RT*_Median_is the median RT over all target shapes. The *LI*_general_ index gives us a measure of the overall strategy employed by each subject across conditions. We then split participants based on whether they used the statistical regularities, using a median split. Following the terminology introduced in previous work (Tarasi, Alamia, and Romei 2025; Tarasi, di Pellegrino, and Romei 2022), we label the two groups ‘Believers’ and ‘Empiricists’, based on whether they rely more on statistical rules (the former) or the sensory information (the latter).

For each group, we performed repeated-measures ANOVAs on the same dependent variables, considering CONDITION as a fixed factor and GROUP as a between-subjects factor.

### Pupillometry

All pupil traces were preprocessed using MATLAB custom scripts. First, we smoothed and down-sampled the data using a moving average with a window size of 100 timepoints and no overlap, thereby reducing the sampling rate from 1000Hz to 10Hz. Next, we identified and removed eye blinks by differentiating the signal and applying a threshold estimated from a linear function of the Standard Deviation (STD) across blocks and subjects. Additionally, we filter the data by removing all datapoints that fall 2 STDs away from the average, before interpolating the missing signal using linear interpolation. Lastly, we segmented the data from −0.2 to 3s around each shape onset and baseline-corrected the data, using a baseline between −0.1s to the shape onset. Since target events are known to elicit a significantly stronger pupil response than non-target events (Privitera et al. 2010), our analysis here is only focused on non-target events (target effect quantification and surprisal analysis is shown in Supplementary materials). In order to compare pupil responses after rare and expected events, we calculated the mean pupil response amplitude between 0 and1s after rare and expected non-target events. Additionally, we analyzed, in the same time window, the peak latency difference between rare and expected non-target events in all conditions. We define latency as the time point at which the pupil reaches its maximum response to both rare and expected events. For these two measures, amplitude and peak latency, we performed repeated-measures ANOVAs, considering CONDITION and SHAPE PROBABILITY as fixed factors.

### Electrophysiology

#### Preprocessing

Custom functions of the EEGlab toolbox (Delorme and Makeig 2004) were used for the EEG pre-processing. We first applied average referencing and removed power-line artefacts using a notch filter (47–53Hz). We removed slow drifts by applying a high-pass filter (2Hz), and cut off higher frequencies with a low-pass filter (60Hz). The data were then down-sampled to 160Hz, and noisy channels were rejected. We then performed Independent Component Analysis (ICA) to identify artifacts in the data and used the ICLabel plugin to automatically detect artifactual ICA components. Specifically, we flagged muscle, eye, and heart components identified with a probability of 80% or more and removed them. After this step, the removed channels were interpolated using superfast spherical interpolation.

#### ERPs quantification and statistics

Average ERPs were computed separately for each participant, electrode and experimental condition. We epoched the data from −1.5 to 2s around the shape onset. We applied a baseline correction from −0.2 to 0s before shape onset. For each condition, ERP responses were divided into target and non-target events, and then, further into rare and expected events (excluding C1, which has no rare or expected events).

Similarly to pupil diameter analysis, since target events are known to elicit a stronger and slower P300 component compared to non-target events (Ninomiya et al. 1998; Squires et al. 1977), our analysis here is only focused on non-target events (target analysis is shown in Supplementary materials). The P300 is usually measured within a time window of 250-500ms after event onset (Polich 2007). Therefore, we measured the P300 amplitude (μV) as the average value over this time window, and P300 latency (ms) as the time from stimulus onset to the point of maximum positive amplitude within the same time window. On both measures, amplitude and latency, we performed repeated-measures ANOVAs with CONDITION and SHAPE PROBABILITY as fixed factors. As we did not expect any lateralization, the analysis was performed on four electrodes along the midline (Oz, Pz, Cz, Fz).

#### Alpha Power analysis

To obtain the power spectra, we created data epochs from −1.5s to 2s around each shape onset in the pre-processed data. For each condition, we calculated the time-frequency response by applying a continuous Morlet wavelet on each event separately and averaging the resulting spectra.

We then averaged over the alpha band (7-13Hz) between 0 to 1s after event onset, and, following previous literature (Tarasi et al. 2025, 2022), we computed the Δα index for each participant and experimental condition as follows:

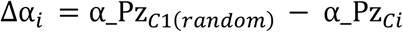

Where α_Pz_C1(random)_and α_Pz_Ci_ represents the average alpha power in electrode Pz in the random condition (C1) and in the other conditions. The Δα provides an index of occipital alpha modulation. To assess the effect of condition on this index, we performed repeated-measures ANOVAs, with CONDITION as a fixed factor. To compare this index with the LI defined above, we performed, for each condition, Kendall’s correlation analysis between the *LI*_cond_and the Δα. Lastly, on the average alpha power of the electrode Pz, we performed repeated-measures ANOVAs, with CONDITION as a fixed factor and, to examine the effect of the *LI*_general_, we added GROUP as a between-subjects factor. We also performed Kendall’s correlation analysis between the *LI*_general_and the average alpha power on electrode Pz across conditions. In Supplementary materials we perform the same analysis on the other midline electrodes Oz, Cz and Fz.

#### Wave quantification

We quantify the presence of TWs and their direction using two different and complementary methods. The first method is based on a 2D Fast Fourier Transformation (2D-FFT) (Alamia and VanRullen 2019) (Figure 1D, upper panel). For each trial, we extracted EEG signals from seven midline electrodes (from posterior to frontal: Oz, POz, Pz, CPz, Cz, FCz, Fz) to form a 2D (electrode-time) map. We applied a 500ms sliding window with a 100ms overlap to each trial and transformed the data using a 2D-FFT. This transform yields a 2D-spectral representation, with temporal frequencies along the horizontal axis and spatial frequencies along the vertical axis. The horizontal midline indicates stationary oscillations with no spatial propagation (i.e., standing waves), while the upper and bottom quadrants reflect FW and BW waves, respectively. We extracted the maximum value across all spatial frequencies from the upper and lower quadrants of the 2D-FFT spectrum to obtain two 1D-spectra over temporal frequencies for both forward and backward waves. For each spatio-temporal map, we normalized both spectra using the mean 1D-FFT over the seven electrodes (Zeng et al. 2024), and we extracted the alpha band. This procedure yields a measure of wave intensity in dB units, as used in previous work (Bailey, Pacheco, and Smillie 2025; Luo and Ester 2025; Pang (庞兆阳) et al. 2020). Eventually, we extracted the average alpha FW and BW waves across all blocks and, on both measures separately, we performed repeated-measures ANOVAs considering CONDITION as a fixed factor. The second method employs a sphere-fitting procedure (Schwenk and Alamia 2025) and was recently introduced for the detection and quantification of planar TWs in scalp-EEG signals (Figure 1D, lower panel). This method was adapted from a previous intracortical recording-based method (Zhang et al. 2018) and was applied here with a fixed Region of Interest (ROI) and Frequency of Interest (FOI). As we didn’t expect any lateralization, we defined a narrow central ROI (N=19 electrodes) extending one electrode to either side from the midline. The FOI comprised the alpha band (7-13Hz) as in the previous analysis. In the first step, the method identifies clusters of true oscillatory activity (as compared to the aperiodic activity), achieved by estimating background noise via an aperiodic fit on the data. Here, since ROI and FOI are fixed, the clusters are defined only in the temporal dimension, as consecutive periods of above-threshold alpha power. A spherical planar wave is then fit to the spatiotemporal phase patterns within each cluster. For each time point, the best fit provides measures of wave direction (WD), spatial frequency and goodness-of-fit. Using these results, we calculated the probability of different states based on WD, considering only fits for which the goodness-of-fit was above a certain threshold (Phase Gradient Directionality, PGD>0.5; for details, see (Schwenk and Alamia 2025; Zhang et al. 2018)). These states are forward, backward, and mediolateral waves within a range of 45° (i.e., ±22.5° from the midline). We computed the probability for each state and performed two analyses on these measures. First, we performed one analysis considering the average FW and BW waves across all blocks, and then conducted repeated-measures ANOVAs, with CONDITION as a fixed factor for FW and BW separately. In order to assess the relationship with overall FW and BW waves and the participant’s strategy used to perform the task, we also estimated the *LI*_general_group effect, by adding the factor GROUP as a between-subject factor in the repeated-measures ANOVA. Then we performed Kendall’s correlation analysis between BW and FW waves and the *LI*_general_and between FW and BW waves and Δα averaged across conditions. In addition, to investigated whether TW are modulated by event onset, we performed analyses on the segmented epochs from −1.5 to 2s around each shape onset. This analysis was not applied to the 2D-FFT method due to its relatively poor time resolution. We performed a timepoint-by-timepoint multiway ANOVA with FDR correction to test for the effects of multiple factors (CONDITION and SHAPE PROBABILITY or TARGET).

### Statistical analysis

The timepoint-by-timepoint ANOVA with FDR correction for the TWs was performed in MATLAB. The CBP analysis performed for the time-frequency data was performed using the FieldTrip toolbox (Oostenveld et al. 2011). The remaining statistical analyses were performed in JASP (Love et al. 2019). For all measures on which we performed repeated-measures ANOVA or Kendall’s correlation, we used both classical and Bayesian methods, and we reported both p-values, as a measure of the significance of the effect, and Bayes Factor, as a measure of its effect size. Bayesian analyses provide Bayes Factors (BFs), which quantify the ratio between the null and alternative hypotheses. In this study, we report BFs as BF10, where larger values provide stronger evidence in favor of the alternative hypothesis.

## Results

### Behavioral

The repeated-measures ANOVA analysis of the behavioral performances (Figure 2A) showed a significant CONDITION effect for False Alarms, (BF10>3 and p-value<0.05), but no significant effect of CONDITION for both Miss and Hit (BF10=0.3 and p-value>>0.05). We found a significant effect of CONDITION also for the perceived predictability scores (questionnaire ratings) (BF10=10^22^ and p-value<0.001), with higher rates for more predictable sequences. Regarding the RT (Figure 2B, right-most panel), we found a very strong effect of SHAPE PROBABILITY and its interaction with CONDITION (BF10=∞ and p-value<0.001 for both), and a weaker effect of CONDITION (BF10=1.12 and p-value<0.001). Overall, these results suggest that the regularities influenced participants’ behavior and that they were aware of them, as indicated by the RTs and questionnaire ratings, respectively.

**Figure 2.**
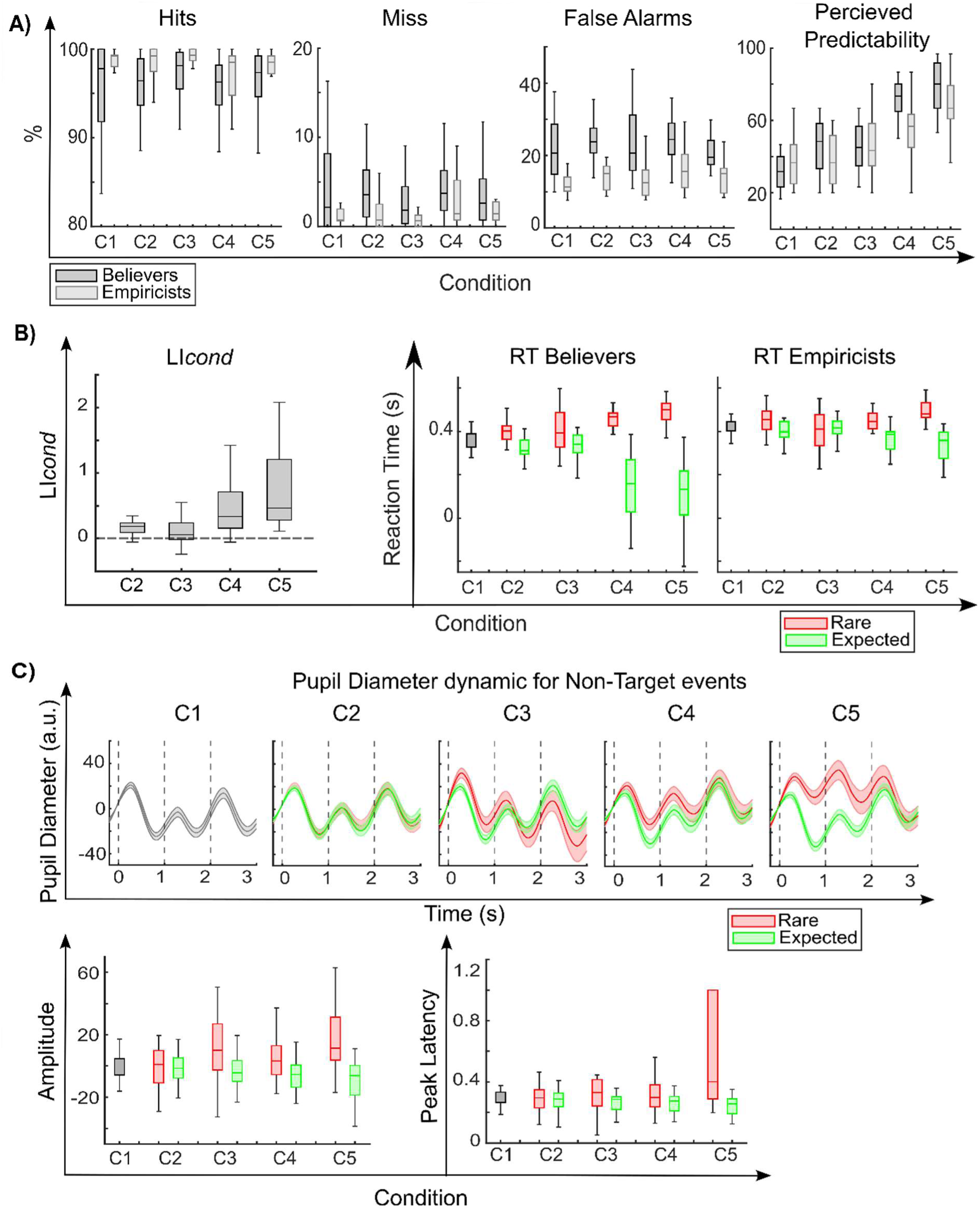
A) Behavioral results. From left to right panel: percentage of hits, miss and false alarms, perceived sequence predictability for the two groups of subjects. 2) RT results. On the left the *LI*_cond_as a function of condition, and on the right the average RT to target events as a function of condition for the two groups of subjects (Believers on the left and Empiricists on the right). C) Pupillary responses. Upper panel: pupil diameter dynamic for Non-Target events: pupillary response as a function of time after shape onset (at t=0) for rare and expected transitions (respectively in red and green) for each condition. Lower panel: average pupil diameter amplitude between 0 and 1s for non-target events as a function of experimental condition; pupil peak latency as a function of experimental condition for non-target events. The box chart shows the median, the lower and upper quartiles, and the minimum and maximum values that are not outliers (computed using the interquartile range). In the pupil diameter dynamic graphs, the shaded area represents the standard error of the mean. C1 = random condition [0.33 0.33 0.33], C2 = condition [0.60 0.20 0.20], C3 = condition [0.475 0.475 0.05], C4 = condition [0.80 0.10 0.10], C5 = condition [0.90 0.05 0.05]

To further characterize participants’ strategies, we used reaction times to define an LI to assess whether they used statistical regularities (Figure 2B, left panel). We found that this index increased with predictability when computed separately per condition (*LI*_cond_, see methods for details), suggesting that participants rely more on the regularities as the sequence predictability increases (repeated-measures ANOVA, effect of CONDITION: BF10=10^+8^and p-value<0.001).

Next, we split participants into two groups based on the LI computed over all conditions (*LI*_general_), and we compared behavioral outcomes between the two groups. As expected, the behavioral results (Figure 2A and 2B) show significant differences between the two groups. Regarding the percentage of miss hit and false alarms, repeated-measures ANOVA showed a strong effect of GROUP (BF10>3 and p-value<0.05), but no significant interaction between GROUP and CONDITION (BF10<0.3 and p-value>0.05). In other words, the ‘Believer’ group has a significantly higher miss and false alarms and fewer hits rate than the ‘Empiricists’, independent of the condition, confirming a much more liberal approach for the ‘Believers’. Concerning the perceived predictability rating, we found a strong effect of CONDITION and GROUP-CONDITION interaction (BF10>3 and p-value<0.001), but no effect of GROUP (BF10<0.3 and p-value>0.05), suggesting that both groups were aware of the regularity level of the experimental condition. Finally, for the RT, we found a strong effect of SHAPE PROBABILITY, GROUP, and their interaction (all BF10>3 and p-value<0.001), revealing that both groups had a decrease in RT for expected targets, proportional to the target’s expectancy, but this decrease was larger for the ‘Believers’ group. These results confirm that the two groups reflect opposite strategies: one more conservative, relying mostly on sensory evidence while disregarding predictability, and one more liberal, using statistics to anticipate the target’s occurrence.

### Pupillometry

Figure 2C shows the difference in the dynamics of pupillary response for rare and expected non-target events. The effect of shape probability, i.e., rare or expected events, is calculated by determining the mean pupil diameter amplitude and peak latency after rare and expected non-target events between 0 and 1s after event onset. As shown in Figure 2C (lower panel), for the amplitude, repeated-measures ANOVAs indicate a weak but significant effect of CONDITION (0.3<BF10<3 and p-value<0.001) and a strong effect of SHAPE PROBABILITY and interaction (BF10>>3 and p-value<0.001). For peak latency, instead, ANOVA analysis indicate a strong effect of CONDITION, SHAPE PROBABILITY, and interaction (BF10>>3 and p-value<0.05). All in all, these results show that pupillary responses are larger and slower for rare events, depending on the condition. These results confirm that participants learn the statistical regularities and are in line with previous studies (Alamia et al. 2019; Schwiedrzik and Sudmann 2020; Silvestrin, Penny, and FitzGerald 2021; Kong, Chen, and Jia 2025).

### Evoked Response Analysis

Regarding the ERP analysis (Figure 3A), we examined differences in responses to rare and expected non-target events. Notably, ERPs’ late positive components, in particular the P300, have been associated with the processing of unexpected events (Nieuwenhuis, Aston-Jones, and Cohen 2005; Polich 2007). In particular, surprising events elicit a larger P300 component (Donchin 1981; Duncan-Johnson and Donchin 1977; Kolossa, Kopp, and Fingscheidt 2015; Kopp et al. 2016; Mars et al. 2008; Nieuwenhuis et al. 2005; Polich 2007; Seer et al. 2016). We therefore tested whether in our design the amplitude and latency of the P300 component are affected by shape probability and condition. Since we did not expect any laterality, we focus the analysis on four midline electrodes (Oz, Pz, Cz, Fz), while also inspecting the topographic distribution of the effect.

**Figure 3.**
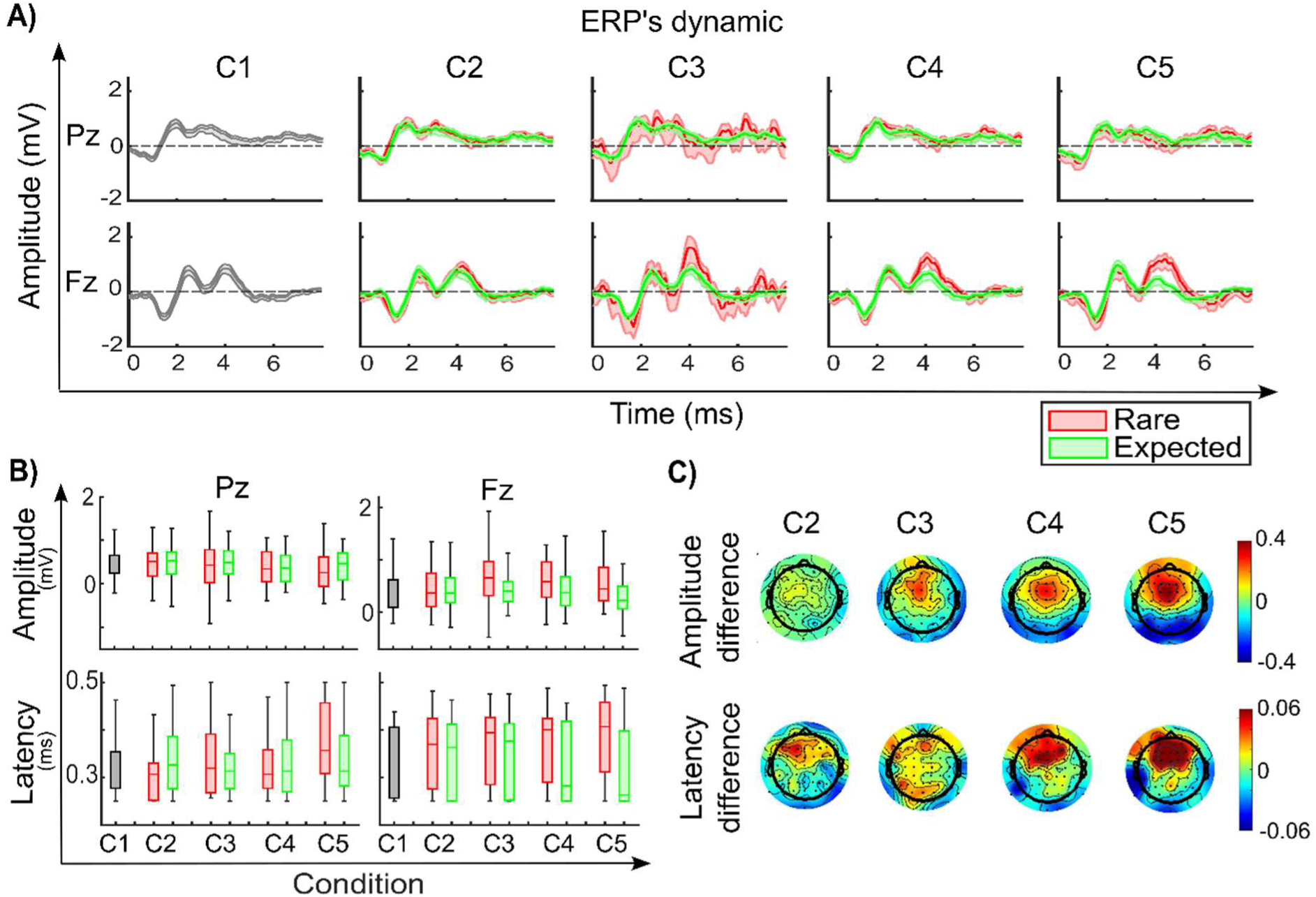
ERP following non-target events A) ERP dynamic. ERP responses to rare and expected events (in red and green, respectively) at electrodes Pz and Fz across the five conditions. B) P300 statistic. On the upper panels, the average P300 amplitude (calculated as the average amplitude between 0.25 and 0.5s after shape onset), on the lower panels the P300 latency (calculated as the time of maximum amplitude between 0.25 and 0.5s after shape onset), both for rare and expected events as a function of condition for electrodes Pz and Fz. C) P300 topography. Topography, for condition two to five, of the difference between the rare and expected P300 amplitude (first line) and latency (second line). The box charts show the median, the lower and upper quartiles, and the minimum and maximum values that are not outliers (computed using the interquartile range). C1 = random condition [0.33 0.33 0.33], C2 = condition [0.60 0.20 0.20], C3 = condition [0.475 0.475 0.05], C4 = condition [0.80 0.10 0.10], C5 = condition [0.90 0.05 0.05].

Amplitude analysis (Figure 3B, upper panel) revealed that, or electrode Oz and Fz there is no effect of CONDITION (BF10<0.3 and p-value>0.05), and a strong effect of SHAPE PROBABILITY and factor interaction (BF10>>3 and p-value<0.001); for Pz there is a weak effect of CONDITION (BF10=1.14 and p-value<0.05), and no effect of SHAPE PROBABILITY or factor interaction (BF10<0.3 and p-value>0.05); for electrode Cz there is a strong effect of both CONDITION, SHAPE PROBABILITY and their interaction (BF10>>3 and p-value<0.001). The topography of the amplitude difference between rare and expected events (Figure 3C, upper panel) shows a stronger difference in centro-frontal areas, which increases with the condition. This suggests that the fronto-central P300 component increased following a rare event compared to an expected one, and that this difference is larger with rarer events, suggesting the involvement of frontal areas in detecting violations in the statistical regularities (Karuza et al. 2013; Schapiro and Turk-Browne 2015; Brylka et al. 2025; Turner et al. 2004).

Peak latency analysis (Figure 3B, lower panel) showed that, for electrode Oz and Pz there is no effect of CONDITION, SHAPE PROBABILITY and their interaction (BF10<0.3 and p-value>0.05); for electrode Cz there is a weak effect of SHAPE PROBABILITY (BF10=1.2 and p-value<0.05), and no effect of CONDITION or factor interaction (BF10<0.3 and p-value>0.05); for electrode Fz there is no effect of CONDITION (BF10<0.3 and p-value>0.05), and a strong effect of SHAPE PROBABILITY and factor interaction (BF10>>3 and p-value<0.001). We investigated the topography of the latency difference (Figure 3C, lower panel) between rare and expected events and observed a stronger lag in frontal areas that increases with the condition, indicating that fronto-central P300 component is slower after a rare event then after an expected one. Also, latency is not affected by sequence predictability. Still, in frontal electrodes, we observed a difference in latency between rare and expected events for more predictable conditions (C4 and C5).

### Alpha-band Oscillatory Activity

#### Alpha-band Power

Regarding the power analysis, we first examined the overall occipital alpha-power difference between the conditions. According to previous literature (Tarasi et al. 2025, 2022), parieto-occipital alpha power is modulated by participants’ task strategy, leading to decreased power for a prior-driven strategy and increased power for stimulus-driven strategy. As we hypothesized that participants are likely to switch their target detection strategy from stimulus-driven to prior-driven as the sequence becomes more predictable, we investigated whether sequence predictability affects occipital alpha power, by calculating the Δα index (see Methods for details). This index shows the extent to which each participant undergoes occipital alpha modulation in each condition relative to the random (stimulus-driven) condition. The bigger Δα is, the more the subject undergoes occipital alpha power suppression. Repeated-measures ANOVA showed that for Δα (Figure 4A) there is an effect of CONDITION (BF10=3.6 and p-value<0.05): Δα increases with sequence predictability, suggesting, in accordance with previous literature (Tarasi et al. 2025, 2022), that participants undergo a strategy change with the different conditions that is reflected by occipital alpha suppression. We also correlated the Δα index with the behavioral *LI*_cond_index, since the latter suggested that participants shift strategy going toward a prior-driven with sequence predictability, and that previous literature (Tarasi et al. 2025, 2022) indicated that participants who use more regularities (i.e. higher *LI*_cond_) also have larger occipital alpha power modulation (higher Δα). We found evidence in favor of a lack of correlation between the two measures (BF10<0.3 and p-value>0.05), suggesting that they may reflect different mechanisms.

**Figure 4.**
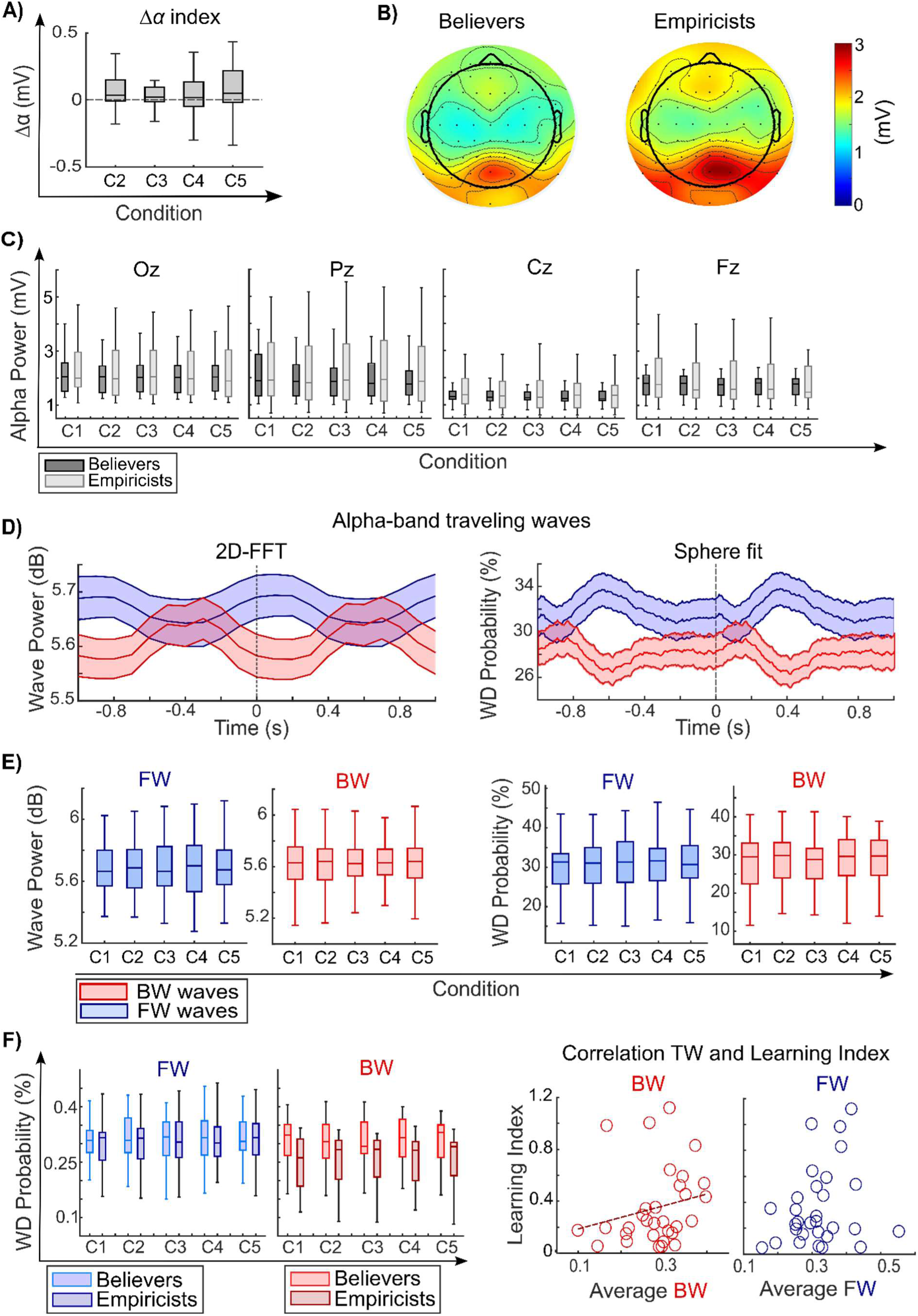
Alpha-band analysis. A) Occipital Δα index. The panel shows the Δα index, as a function of sequence predictability. B) Topography of alpha power. Average alpha power across experimental condition for ‘Believers’ (left panel) and ‘Empiricists’ (right panel). C) Alpha power across conditions. The panels show the average alpha power for the two participant’s groups (‘Believers’ in dark grey and ‘Empiricists’ in light gray), as a function of sequence predictability, for the electrode Oz, Pz, Cz and Fz. D) Wave dynamics for FW and BW waves. Dynamics of FW and BW waves (respectively represented in blue and red) around the event onset. The left panels display the results for the 2D-FFT method in decibels, whereas the right panels show the results for the Sphere fit method as Wave Direction probability. E) Overall FW and BW wave strength. Average strength of FW and BW waves across all blocks, as a function of sequence predictability. F) Alpha TWs and LI results. Left panel shows alpha band TWs results: overall FWs and BWs for the two groups as a function of condition. Right panel shows correlation between overall BW and FW waves, averaged across condition with Learning Index. The scattered line represents the least-squares fit. In all panels, the shaded area represents the standard error of the mean. The box charts show the median, the lower and upper quartiles, and the minimum and maximum values that are not outliers (computed using the interquartile range). C1 = random condition [0.33 0.33 0.33], C2 = condition [0.60 0.20 0.20], C3 = condition [0.475 0.475 0.05], C4 = condition [0.80 0.10 0.10], C5 = condition [0.90 0.05 0.05].

We then investigated the difference in alpha power between the two participants groups. Figure 4C shows the average alpha power of the four midline electrodes for the two groups. Repeated-measures ANOVA showed that for electrode Oz there is a weak effect of CONDITION (BF10=2.5 and p-value<0.05); for electrode Pz, Cz and Fz there is a strong effect of CONDITION (BF10>>3 and p-value<0.05). This result indicates a reduction in overall alpha power under more predictable conditions, especially in central areas, in line with previous literature (Tarasi et al. 2025, 2022) that showed that prior-driven conditions result in desynchronization of alpha power, whereas higher alpha power is observed in stimulus-driven conditions. Furthermore, for all four electrodes there is an inconclusive effect of GROUP (0.3<BF10>3 and p-value<0.05) and no interaction CONDITION-GROUP for all electrodes (BF10<0.3 and p-value>0.05). Despite the inconclusive results, the alpha power topography shown in Figure 4B (averaged across condition) suggests a trend in line with previous results, with the group of the ‘Believers’ having lower alpha power compared to the ‘Empiricists’. Furthermore, the correlation between *LI*_general_ and alpha power across conditions for four midline electrodes revealed no effect (Kendall correlations, BF10<0.3 and p-value>0.05).

#### Traveling waves

Concerning the alpha-band traveling wave analysis, we focused on waves propagating from the occipital to the frontal regions (FW) and in the opposite direction (BW). We first investigate the dynamics around event onset (Figure 4D), regardless of condition or target, using the two detection methods: the 2D-FFT and the Sphere fit (see Methods for details). As discussed in previous work (Schwenk and Alamia 2025), the two methods differ significantly in several aspects: the temporal resolution is much higher for the Sphere fit, as the 2D-FFT considers overlapping time-windows of 500ms and may suffer from smearing effects (which explains the increase in FW before stimulus onset). On the other hand, the Sphere fitting considers only trials and time points with high spectral power, whereas the 2D-FFT relies on the average across all trials. 2D-FFT reveals an increase in FW waves before the event onset, and an increase in BW waves around shape onset, while the Sphere fit method revealed an increase in FW waves around 300ms after the event onset and in BW waves immediately after the event onset.

Next, we tested whether FW and BW waves are modulated by the predictability condition across all blocks (regardless of type and event onset). In particular, we hypothesized that participants would rely more on sensory evidence rather than predictions in more unpredictable conditions (i.e., C1), and this would be reflected in an overall increase of FW waves; conversely, in more predictable conditions (i.e., C5), participants would rely more on the regularities, thus having an overall increase in BW waves. As shown in the Figure 4E, for both FW and BW and using both TW detection methods, we found a convincing lack of effect for the factor CONDITION (BF10<0.3 and p-value>0.05), thus providing evidence against the hypothesis that alpha-band TW reflect the learning of statistical regularities.

We then investigated whether TW are modulated by event onset. Given the relatively poor temporal resolution of the 2D-FFT method, we focused this analysis on the Sphere fit method. We first investigate changes in FW and BW waves around shape onset (regardless of the type of event) as a function of the condition. FW waves, baseline corrected between −0.3 and 0s before event onset, revealed no difference between conditions, as confirmed by a timepoint-by-timepoint multiway ANOVA considering the effect of CONDITION and including FDR correction (for all timepoints, p-value>>0.05). Similar results were obtained for the BW waves (all p-values>>0.05), which we analyzed with a similar approach but without baseline correction. These results provide clear evidence against the hypothesis that alpha band FW and BW waves are modulated by sequence predictability in relation to event onset.

We then investigated whether the event type (target, non-target, rare or expected) modulates wave strength. We observed no difference in baseline-corrected FW waves between rare and expected events as well as between target and no target (all p-values>>0.05 in a timepoint-by-timepoint multiway ANOVA with CONDITION and TARGET or SHAPE PROBABILITY as factors). Interestingly, and contrary to previous hypotheses (Alamia et al. 2025; Alamia and VanRullen 2019; Friston 2019; Mohanta et al. 2024), this result suggests that FW waves may not reflect prediction errors during statistical learning, nor do they relate to the saliency of the target onset. Concerning the BW waves, our analysis did not detect any difference between expected target and non-target conditions, as confirmed by a timepoint-by-timepoint multuway ANOVA with CONDITION and TARGET as factors (all FDR-corrected p-values>>0.05). This last analysis supports the hypothesis that BW waves are not modulated by sequence predictability and target prediction.

#### Traveling waves and Learning Index

To characterize participants’ strategies relationship with TWs, we performed the same group-level analysis on the TW probabilities (as derived from the Sphere fit method). As shown in Figure 4F (left panels), we found no effect of both CONDITION and GROUP for the FW waves, nor their interaction (BF10<0.3 or 0.3<BF10>3 and p-value>0.05). For BW waves, the analysis revealed no effect of either CONDITION and their interaction (BF10<0.3 and p-value>0.05), but a weak effect of GROUP (BF10=1.6 and p-value<0.05), indicating a higher amount of BW waves in the ‘Believers’ group. Interestingly, a Kendall’s correlation between *LI*_general_ and traveling waves averaged across condition (Figure 4F, right panels) revealed a correlation for BW waves (BF10=2.2 and a p-value<0.05), but not for FW waves (BF10<0.3 and p-value>0.05).

## Discussion

The goal of the present study was to explore the link between alpha-band oscillations and predictive processes using a visual statistical learning paradigm. Our results show that regularities influenced participants’ behavior both in their RT and awareness of the sequence predictability (higher perceived predictability ratings were assigned to more predictable sequences). Importantly, we show that physiological measures, including pupil size, ERP responses, and spectral power, were also modulated by different levels of regularity. Lastly, our results provide evidence against the hypothesis that alpha-band TWs broadly reflect predictive processes, as alpha waves were not modulated by sequence predictability or by rare events. Importantly, however, we did find a modulation of overall BW waves in relation to the participant’s strategy. Participants who used the regularities more had a higher occurrence of BW waves compared to those who relied more on the stimuli, highlighting a possible role of these waves in shaping cognitive strategy (Tarasi et al. 2025).

Our behavioral results are in line with previous work (Chun and Jiang 1998, 1999; Kim et al. 2009; Jost et al. 2011; Tóth et al. 2017; Zhao et al. 2024) showing that the regularities induced by SL paradigms improve target detection timing by allowing participants to predict and anticipate target occurrence. It is well known that brain arousal affects pupil size, and this effect is thought to reflect increased sympathetic activity (Bradley et al. 2008). One way to induce arousal is surprisal, i.e., the degree of an observation’s unexpectedness. In particular, pupil dilation has been shown to reflect conscious and unconscious surprise, with an inverse relationship between stimulus probability and pupil size (Raisig et al. 2010; Preuschoff’t Hart, and Einhäuser 2011; Nassar et al. 2012; Damsma and van Rijn 2017; Alamia et al. 2019; Schwiedrzik and Sudmann 2020; Silvestrin et al. 2021; Kong et al. 2025). In line with these findings, our data show slower, more pronounced pupil dilation following novel or surprising information than following expected information, with an effect increasing with the rarity of the event. We also found a strong modulation by both event surprisal and sequence predictability in the ERP analysis of the late component, P300. The P300 has previously been shown to be enhanced by surprising events (Donchin 1981; Duncan-Johnson and Donchin 1977; Kolossa et al. 2015; Kopp et al. 2016; Mars et al. 2008; Seer et al. 2016), and has been related to context-updating mechanisms (Donchin 1981; Duncan-Johnson and Donchin 1977; Polich 2007; Polich and Comerchero 2003), and to the Bayesian brain hypothesis (Kolossa et al. 2015; Kopp et al. 2016; Mars et al. 2008; Seer et al. 2016). In this framework, the P300 would be interpreted as a result of prediction error transmission. In line with this hypothesis, we found that for non-target events, the amplitude of the fronto-central P300 component is higher, and its latency slower, following unexpected shape transitions as compared to expected ones, and this effect increases as the predictability of the stream of shapes is higher (i.e., the more surprising the unexpected event is). All in all, along with the behavioral results, the pupillary and ERP results provide robust evidence that participants learned the statistical regularities embedded in the sequence.

Previous work has proposed the hypothesis that FW and BW alpha-band traveling waves relate to predictive processes (Alamia and VanRullen 2019; Friston 2019). Computational models of cortico-cortical interactions (Alamia and VanRullen 2019; Schwenk and Alamia 2024) relate alpha-band FW waves to sensory information and PEs, while BW waves carry top-down predictions. Recent experimental work seems to provide indirect evidence in favor of this interpretation. In particular, supporting the theoretical framework that suggests modulation of precision-weighted prediction errors in specific neuroatypical populations (Friston et al. 2016; Sterzer et al. 2018), experimental work found altered top-down and bottom-up traveling waves in schizophrenia patients (Alamia et al. 2025) and in autistic populations (Alamia et al. 2026). Similarly, psychedelic drugs have been proposed to relax the precision of high-level priors or beliefs, eliciting an increase of bottom-up, prediction error-related activity (Carhart-Harris and Friston 2019). Experimental results analyzing alpha-band traveling waves in healthy subjects during psychedelics intake provided evidence corroborating this interpretation (Alamia et al. 2020). All in all, these results indirectly suggest that FW and BW waves could be modulated by the precision of prior and prediction errors. In this work, we investigated this hypothesis for the first time in scalp EEG recordings using a visual statistical learning paradigm. Our paradigm manipulated shape-sequence predictability to induce expectations and prediction errors. We hypothesized that (1) sequence predictability would modulate overall FW and BW activity across blocks; (2) FW and BW would vary with event onset and type (rare vs. expected; target vs. non-target). Specifically, we predicted stronger FW responses to salient events—especially rare targets—consistent with FW reflecting prediction errors. Finally, if pre-stimulus BW carries prior information, we expected higher BW before target events (given their task relevance) than non-targets, with this increase scaling with event expectedness. Contrary to our hypotheses, although behavioral and physiological data confirmed that participants learned the statistical regularities, our findings do not support the idea that alpha-band TWs reflect predictive processes in visual statistical learning. FW and BW waves were not modulated by sequence predictability, either overall or at event onset. FW did not index prediction errors or target salience, and BW was unaffected by predictability or target expectation. This may indicate that our paradigm did not induce sufficiently strong expectations to modulate wave strength, or that any modulation was too subtle to detect. It is worth noting that experimental work with intracortical recordings found a relationship between the phase of alpha-band oscillations and the predictability of cues and targets (Mohanta et al. 2024), suggesting that different experimental designs or recordings with a higher signal-to-noise ratio could better delineate the validity of this hypothesis. One main limitation of prior work using scalp-EEG recordings is that earlier results were mainly obtained during closed-eyes resting states (Alamia et al. 2020, 2025), i.e., in the absence of visual stimulation. It is possible that the high-amplitude visual responses evoked by repeated stimulation in our paradigm masked any modulation of BW and FW waves induced by the predictability. A different design may provide a means to better disentangle visual responses from higher-order predictive processes. Interestingly, however, the only effect we found for TWs was between groups of subjects (with ‘Believers’, i.e., subjects who relied more on sequence regularities and exhibited more BW waves). Together with the results from resting-state analyses, this could suggest that modulation of TWs reflects global network states on a longer time-scale (e.g., cognitive strategy, arousal) rather than local processing dynamics at a time-scale of a few oscillatory cycles, as in the event response. This interpretation is also in line with the modulation of TWs by attentional state (Alamia et al. 2023). Future studies may rely more strongly on comparisons between groups of subjects or induce modulations at longer time scales.

It is important to note that a precise understanding of the neural correlates of macroscopic traveling waves is still being developed. It has been shown that two spatially discrete dipole sources at the scalp level (Zhigalov and Jensen 2023) or multiple discrete clusters of time-lagged activity (Orsher et al. 2024) could explain the observed cortical traveling wave measured by EEG and MEG, consistent with the idea that traveling waves at the scalp level result from the summation of neuronal activity generated by phase-lagged dipoles. These accounts, all assuming inter-areal communication as the correlate of macroscale waves, formed the basis for our hypotheses about a possible association between TWs and top-down modulation. However, recent studies have shown that mesoscopic TWs (i.e., within a single cortical area) can also give rise to global waves in EEG/MEG (Orczyk and Kajikawa 2022; Petras, Grabot, and Dugué 2025). This account is supported by recordings at the cortical level showing that the evoked response in early visual areas (V1/V2) propagates locally as a traveling wave (Muller et al. 2014, 2018b). A potential influence of these mesoscale components provides an alternative explanation why we did not find a direct modulation of TWs by expectation.

Importantly, the difference we observed in the total amount of BW waves depending on the subject’s task strategy matches that observed previously (Tarasi et al. 2025). However, unlike that study, we did not find the reverse effect for FW waves. This discrepancy is likely due to differences in the task between the two experiments: their study used a target detection paradigm with high visual demand, whereas in our task, the target simply had to be reported. Since, according to our hypothesis, FW waves reflect visual sampling and processing, a detection task employing a difficult-to-perceive target could elicit a higher number of FW waves than an easily detectable target.

Remarkably, we replicate the modulation of occipital alpha amplitude following a strategy shift, showing increased alpha suppression in conditions where participants rely more on prior expectations compared to stimulus-driven conditions. We also measured a behavioral marker of the participant’s strategy change across conditions (the *LI*_cond_ index), indicating a shift toward a prior-driven strategy for more predictable conditions. These two measures provide both behavioral and physiological evidence of the strategy change induced by sequence predictability, but, interestingly, we found that these two indices did not correlate, suggesting that they may reflect different mechanisms. Unlike Tarasi and colleagues, who report that a bias shift toward prior-driven strategy is accompanied by occipital alpha suppression, an increase in BW waves, and a decrease in FW waves, in our experimental design, the strategy changes and alpha suppression were not accompanied by a modulation of overall FW and BW waves. However, in line with their study, we observed a relationship between BW waves and the overall participant’s strategy, suggesting that the magnitude of alpha-band BW traveling waves relates to the differences in predictive styles: prior-driven individuals (‘Believers’), showed an increase in BW waves compared to sensory-driven individuals (‘Empiricists’). Revisiting the predictive account of traveling waves, we could hypothesize that BW waves reflect top-down processes by modulating the alpha power in lower regions, which is assumed to integrate prior information and modulate decision-making outcomes (Kloosterman et al. 2019; Tarasi et al. 2022). Our results only partially support this interpretation, as we observed only a trend in the difference in occipital alpha power between the two groups. This suggests that, in our experiment, BW traveling waves play a more global role in mediating the integration of prior expectations, but that, unlike previous literature, this does not occur through a local modulation of occipital alpha power. A possible explanation for this difference could be differences in the experimental design or in how we defined the two groups: we categorized participants based on their behavioral outcome, while previous studies (38,39) defined it based on differences in occipital alpha power desynchronization. Future work will be needed to further disentangle the relationships between these variables.

## Author Contributions

MP: Conceptualization, Data curation, Investigation; Formal analysis, Methodology, Writing – original draft. JCBS: Methodology, Software, Writing – review & editing. AA: Conceptualization, Funding acquisition, Project administration, Supervision, Writing – review & editing.

## Acknowledgements

This work was funded by the European Union under the European Union’s Horizon 2020 research and innovation program (grant agreements No. 101075930 to AA). The copyright holder for this is of the author(s) only and do not necessarily reflect those of the European Union or the European Research Council (ERC). Neither the European Union nor the granting authority can be held responsible for them.

## Supplementary Material

### Pupillometry

Target events elicited a significantly higher and longer response than non-target events (Figure 1A), lasting approximately 2s and being one order of magnitude higher. To quantify the target effect (Figure 1B), we first calculated the mean amplitude for target and non-target events between 0 and 2s. Classical and Bayesian repeated-measures ANOVA showed no significant effect of CONDITION (0.3 < BF10 > 3 and p-value > 0.05), a strong effect of TARGET (BF10 >> 3 and p-value < 0.001), and no interaction between the two (0.3 < BF10 > 3 and p-value > 0.05).

The effect of shape probability, i.e., rare or expected events, is calculated by determining the mean pupil diameter amplitude and peak latency after rare and expected events, respectively, for non-target events between 0 and 1s and for target events between 0 and 2s. Pupil amplitude analysis, as shown in Figure 1C upper panels, revealed that, for targets there was no significant effect of CONDITION (BF10 < 0.3 and p-value > 0.05), a weak effect of SHAPE PROBABILITY (0.3 < BF10 < 3 and p-value < 0.05), and no interaction between the two factors (0.3 < BF10 > 3 and p-value < 0.05), while for non-target events, both repeated-measures ANOVAs reprt a weak but significant effect of CONDITION (0.3 < BF10 < 3 and p-value < 0.001), a strong effect of SHAPE PROBABILITY and factor interaction (BF10 >> 3 and p-value < 0.001).

Concerning the peak latency analysis (Figure 1C, lower panels), for target events there was no effect of CONDITION and factor interaction (BF10 < 0.3 and p-value > 0.05), and a weak effect of SHAPE PROBABILITY (0.3 < BF10 < 3 and p-value < 0.05). For non-target events, repeated-measures ANOVAs indicate a strong effect of both CONDITION, SHAPE PROBABILITY, and factor interaction (BF10 >> 3 and p-value < 0.05).

All in all, these results show that, when the shape is not a target, pupillary responses are larger and slower for rare events, depending on the condition. When the shape is a target, the response shadows any effect of condition or shape probability.

**Figure 1.**
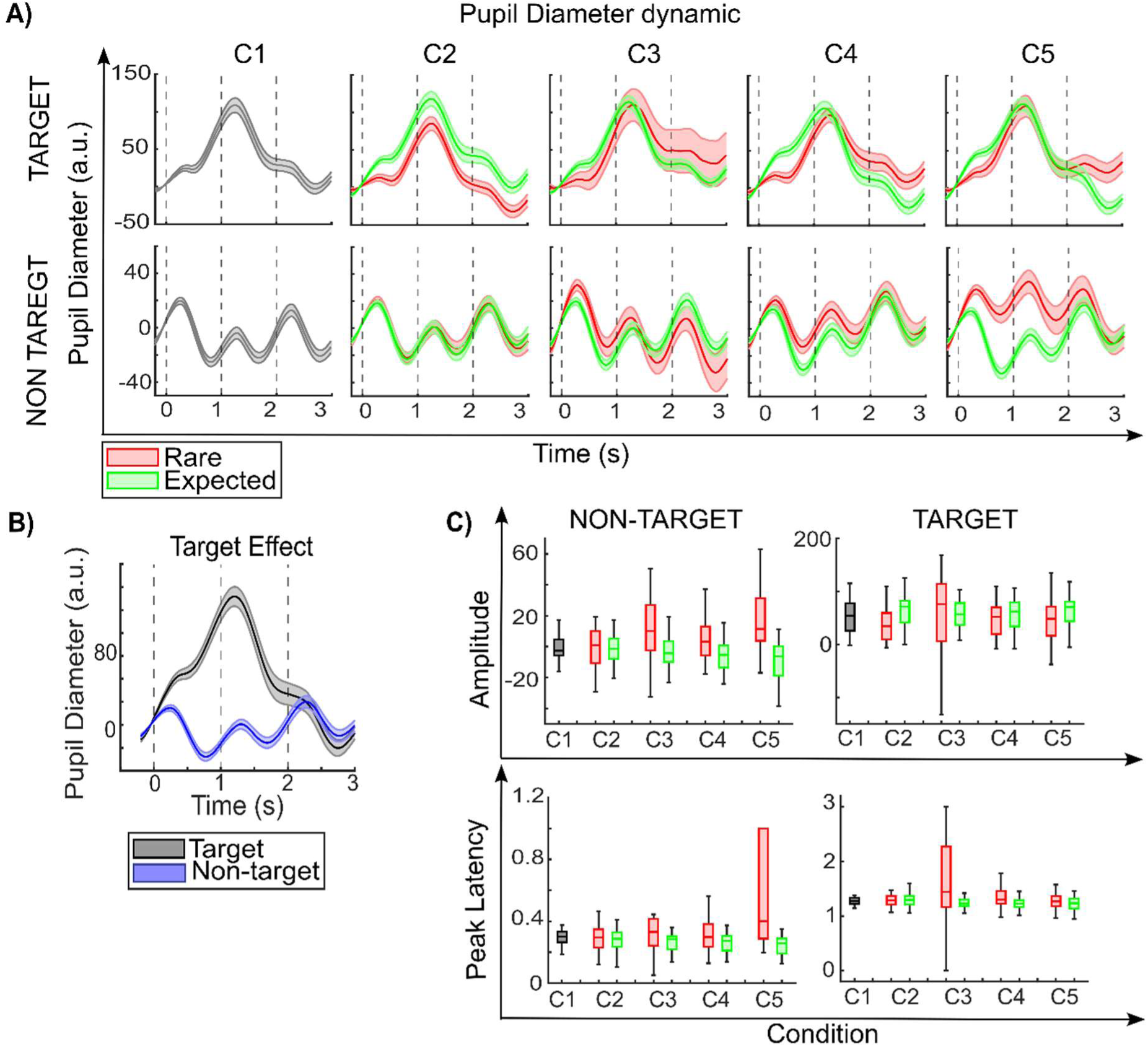
Pupil diameter analysis. A) Pupillary diameter dynamic. Pupil diameter dynamic for target events (upper panels) and non-target events (lower panels): pupillary response as a function of time after shape onset (at t=0) for rare and expected transitions (respectively in red and green) for each condition. B) Target effect. Average target and non-target pupillary responses (in black and blue respectively). C) Amplitude and Peak Latency: average pupil diameter amplitude (upper panes) between 0 and 1s for non-target events (left panels) and between 0 and 2s for target events (right panels) as a function of experimental condition; pupil peak latency (lower panels) between 0 and 1s for non-target events (left panels) and between 0 and 2s for target events (right panels) as a function of experimental condition. The box chart shows the median, the lower and upper quartiles, and the minimum and maximum values that are not outliers (computed using the interquartile range). In the pupil diameter dynamic graphs, the shaded area represents the standard error of the mean. C1 = random condition [0.33 0.33 0.33], C2 = condition [0.60 0.20 0.20], C3 = condition [0.475 0.475 0.05], C4 = condition [0.80 0.10 0.10], C5 = condition [0.90 0.05 0.05]

### Evoked Response Analysis

Hereby we tested whether the amplitude and latency of the P300 component induced after target events are affected by shape probability and condition (Figure 2). As for non-target events, since we did not expect any lateralization, we focus the analysis on midline electrodes (Oz, Pz, Cz, Fz), while also inspecting the topographic distribution of the effect.

We consider the amplitude and latency of the P300 for target events (Figure 2B). Repeated-measures ANOVAs on P300 amplitude (Figure 2B, upper panels) showed that for electrode Oz there is an effect of CONDITION and factor interaction (BF10 > 3 and p-value < 0.001), and no effect of SHAPE PROBABILITY (BF10 < 0.3 and p-value > 0.05); for electrode Pz there is a strong effect of SHAPE PROBABILITY and factor interaction (BF10 >> 3 and p-value < 0.001), and no effect of CONDITION (BF10 < 0.3 and p-value > 0.05); for electrode Cz there is a strong effect of CONDITION, SHAPE PROBABILITY and factor interaction (BF10 >> 3 and p-value < 0.001); for electrode Fz, there is an effect of CONDITION (BF10 > 3 and p-value < 0.05), but no effect of SHAPE PROBABILITY or interaction (BF10 < 0.3 and p-value > 0.05). We investigated the topography of the amplitude difference between rare and expected events (Figure 2C, upper panels): it reveals a stronger difference in centro-parietal areas that increases with the condition. These results show that, in centro-parietal electrodes, the P300 response to expected events is higher than to rare events, increasing with sequence predictability.

Repeated-measures ANOVAs on the P300 latency (Figure 2B, lower panels) revealed that for electrode Oz and Cz there was no effect of CONDITION, SHAPE PROBABILITY and their interaction (BF10 < 0.3 and p-value > 0.05); for electrode Pz there was a strong effect of CONDITION (BF10 = 134.5 and p-value < 0.001), a weak effect of SHAPE PROBABILITY (0.3 < BF10 < 3 and p-value < 0.05), and no factor interaction (BF10 < 0.3 and p-value > 0.05); for electrode Fz there was no effect of CONDITION (0.3 < BF10 > 3 and p-value > 0.05), a strong effect of SHAPE PROBABILITY and factor interaction (BF10 >> 3 and p-value < 0.001). We then investigated the topography of the latency difference between rare and expected events (Figure 2C, lower panels); it reveals a stronger lag in frontal areas for conditions C3 and C5. Altogether, these results show that, in frontal electrodes, the response to rare events is slower compared to expected ones.

**Figure 2.**
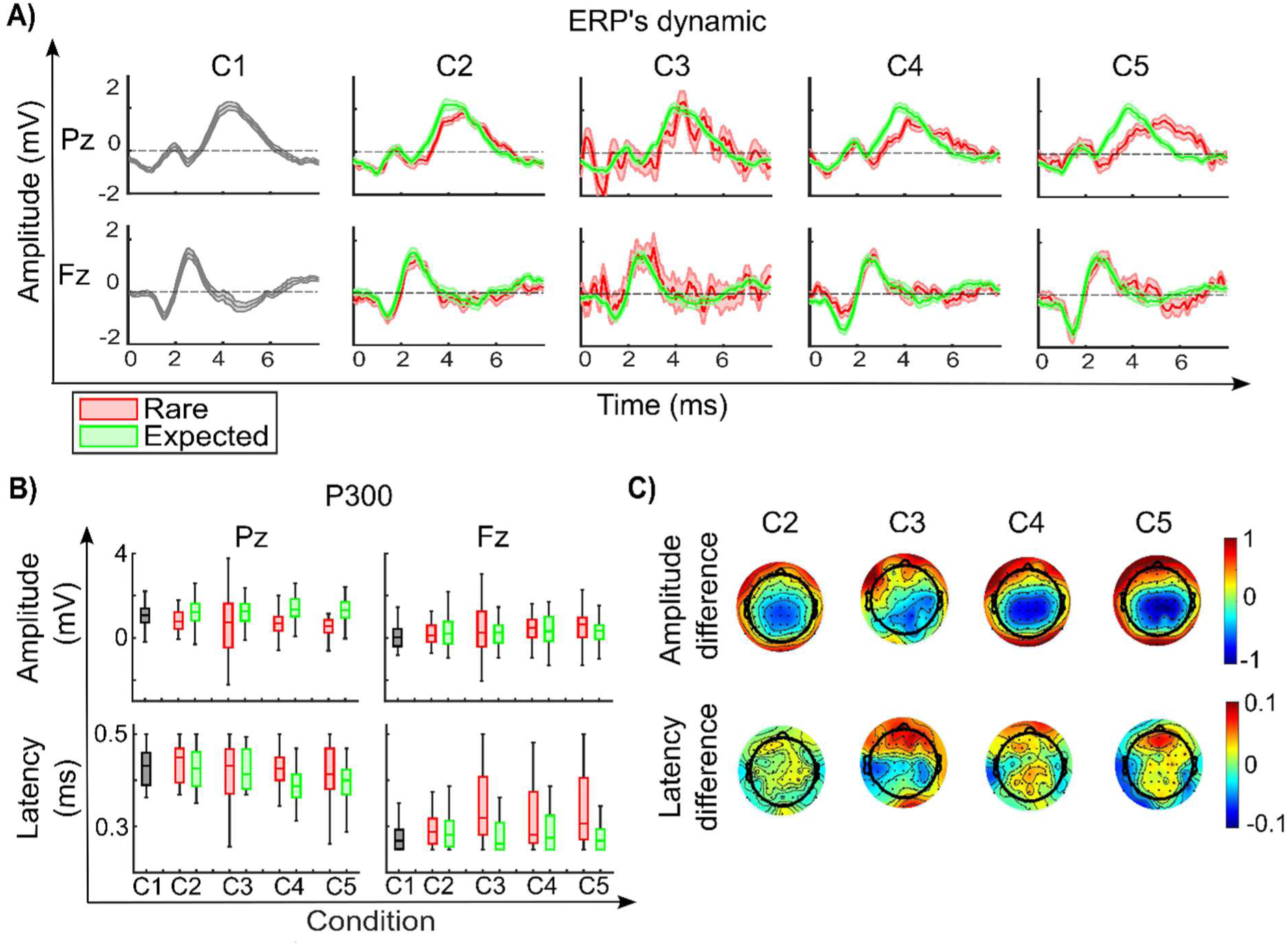
ERP following target events A) ERP dynamic. ERP responses to rare and expected events (in red and green, respectively) at electrodes Pz and Fz across the five conditions. B) P300 statistic. On the upper panels, the average P300 amplitude (calculated as the average amplitude between 0.25 and 0.5s after shape onset) for elctrodes Pz and Fz, on the lower panels the P300 latency (calculated as the time of maximum amplitude between 0.25 and 0.5s after shape onset), both for rare and expected events as a function of condition. C) P300 topography. Topography, for condition two to five, of the difference between the rare and expected P300 amplitude (first line) and latency (second line). The box charts show the median, the lower and upper quartiles, and the minimum and maximum values that are not outliers (computed using the interquartile range). C1 = random condition [0.33 0.33 0.33], C2 = condition [0.60 0.20 0.20], C3 = condition [0.475 0.475 0.05], C4 = condition [0.80 0.10 0.10], C5 = condition [0.90 0.05 0.05].

## Notes

### Competing Interest Statement

The authors have declared no competing interest.

